# Spatiotemporal Maps of Proprioceptive Inputs to the Cervical Spinal Cord During Three-Dimensional Reaching and Grasping

**DOI:** 10.1101/790816

**Authors:** Pierre Kibleur, Shravan R Tata, Nathan Greiner, Sara Conti, Beatrice Barra, Katie Zhuang, Melanie Kaeser, Auke Ijspeert, Marco Capogrosso

## Abstract

Proprioceptive feedback is a critical component of voluntary movement planning and execution. Neuroprosthetic technologies aiming at restoring movement must interact with it to restore accurate motor control. Optimization and design of such technologies depends on the availability of quantitative insights into the neural dynamics of proprioceptive afferents during functional movements. However, recording proprioceptive neural activity during unconstrained movements in clinically relevant animal models presents formidable challenges. In this work, we developed a computational framework to estimate the spatiotemporal patterns of proprioceptive inputs to the cervical spinal cord during three-dimensional arm movements in monkeys. We extended a biomechanical model of the monkey arm with ex-vivo measurements, and combined it with models of mammalian group-Ia, Ib and II afferent fibers. We then used experimental recordings of arm kinematics and muscle activity of two monkeys performing a reaching and grasping task to estimate muscle stretches and forces with computational biomechanics. Finally, we projected the simulated proprioceptive firing rates onto the cervical spinal roots, thus obtaining spatiotemporal maps of spinal proprioceptive inputs during voluntary movements. Estimated maps show complex and markedly distinct patterns of neural activity for each of the fiber populations spanning the spinal cord rostro-caudally. Our results indicate that reproducing the proprioceptive information flow to the cervical spinal cord requires complex spatio-temporal modulation of each spinal root. Our model can support the design of neuroprosthetic technologies as well as in-silico investigations of the primate sensorimotor system.

## I. INTRODUCTION^1^

TRAUMATIC injuries of the central and peripheral nervous system interrupt the bi-directional communication between the brain and the periphery. Neuroprosthetic systems aiming at the recovery of motor function have been mainly focused on the restoration of motor control via direct muscle stimulation [1-3], peripheral nerve stimulation [4-6] and spinal cord stimulation [7-12]. For example, epidural electrical stimulation (EES) [13] of the lumbar spinal cord has shown promising results for the recovery of multi-joint movements in animals [11, 14] and humans [10, 15] with spinal cord injury (SCI). EES engages motoneurons pre-synaptically by directly recruiting large myelinated afferents in the posterior roots [16, 17]. In fact, the stimulation-induced information is processed by spinal circuitry and integrated with residual descending drive and sensory signals to produce coordinated movement [18] [19].

These encouraging clinical results have produced a surge of interest in the application of spinal cord stimulation to the cervical spinal cord to restore also arm and hand movements [12, 20, 21]. However, restoration of voluntary control of arm and hand movements likely requires even finer integration between stimulation signals, descending drive and natural sensory feedback [22]. Unfortunately, electrical stimulation patterns interfere with natural afferent activity [23] leading to impairment of movement execution and conscious perception of proprioception [24]. Therefore, application of EES protocols to the complex control of the upper limb should rely on precise knowledge of cervical sensorimotor circuit dynamics. More generally, any application that aims at restoring limb function [2, 3], or even at the control of external devices [25, 26], might benefit from the restoration of proprioceptive feedback to enhance movement quality and control [23, 27]. In this view, experimental recordings of proprioceptive afferent dynamics are pivotal to future developments in neurotechnologies. Recordings of afferent activity in humans can be performed using microneurography [28, 29], but this technique only allows the recording of single fibers in constrained experimental settings. Alternatively, extracellular recordings of dorsal root ganglion sensory neurons can be obtained in non-primate animal models during functional movements [30, 31]. However, although the latter allows recording multiple fibers simultaneously, it does not readily permit discrimination between the different fiber types, which requires a-priori knowledge of the firing dynamics of each cell type during movement. Moreover, studies addressing the human upper limb sensory dynamics require more pertinent animal models such as non-human primates, in which similar invasive recordings during unconstrained functional movements still present formidable challenges. Here we sought to combine experimental recordings of kinematics and muscle activity in monkeys with a biomechanical model of the primate arm to produce in-silico estimates and characterize the firing rates of proprioceptive fiber ensembles during arm movements. Using OpenSim [33], we extended and scaled the biomechanical model of the *Macaca Mulatta* upper limb developed by Chan and Moran [32], with dedicated ex-vivo measurements, to the size and functional parameters of *Macaca Fascicularis*. We then trained two monkeys to reach and grasp a spherical object while recording arm joint kinematics and electromyograms (EMGs) of the principal arm and hand muscles. We validated this biomechanical model by comparing simulated kinematics and muscle activity with experimental recordings, and used the model to extract muscle stretches and tendon elongation parameters. Next, we fed simulated muscle and tendon states to empirical models of group Ia, Ib and II proprioceptive afferents [34, 35]. Finally, we projected the simulated activity of each of the fiber ensembles onto the spinal segments hosting their homonymous motor pools, thus obtaining spatiotemporal maps of the proprioceptive input to the cervical spinal cord during movement.

## II. METHODS

The computational framework to estimate the firing dynamics of proprioceptive sensory afferents of the upper limb in non-human primates is presented in Fig. 1. It consists of a biomechanical model of the primate’s right arm, fine-tuned to the muscle mechanical properties and anatomy of *Macaca Fascicularis*, and in a mathematical model linking muscle and tendon stretches to firing rates of group Ia, Ib and II afferent fibers. It is complemented with an experimental dataset of the three-dimensional arm joint kinematics and muscle activity of *Macaca Fascicularis* during reaching and grasping movements, and a method to project the afferent activity onto the cervical spinal segments.

**Fig. 1:**
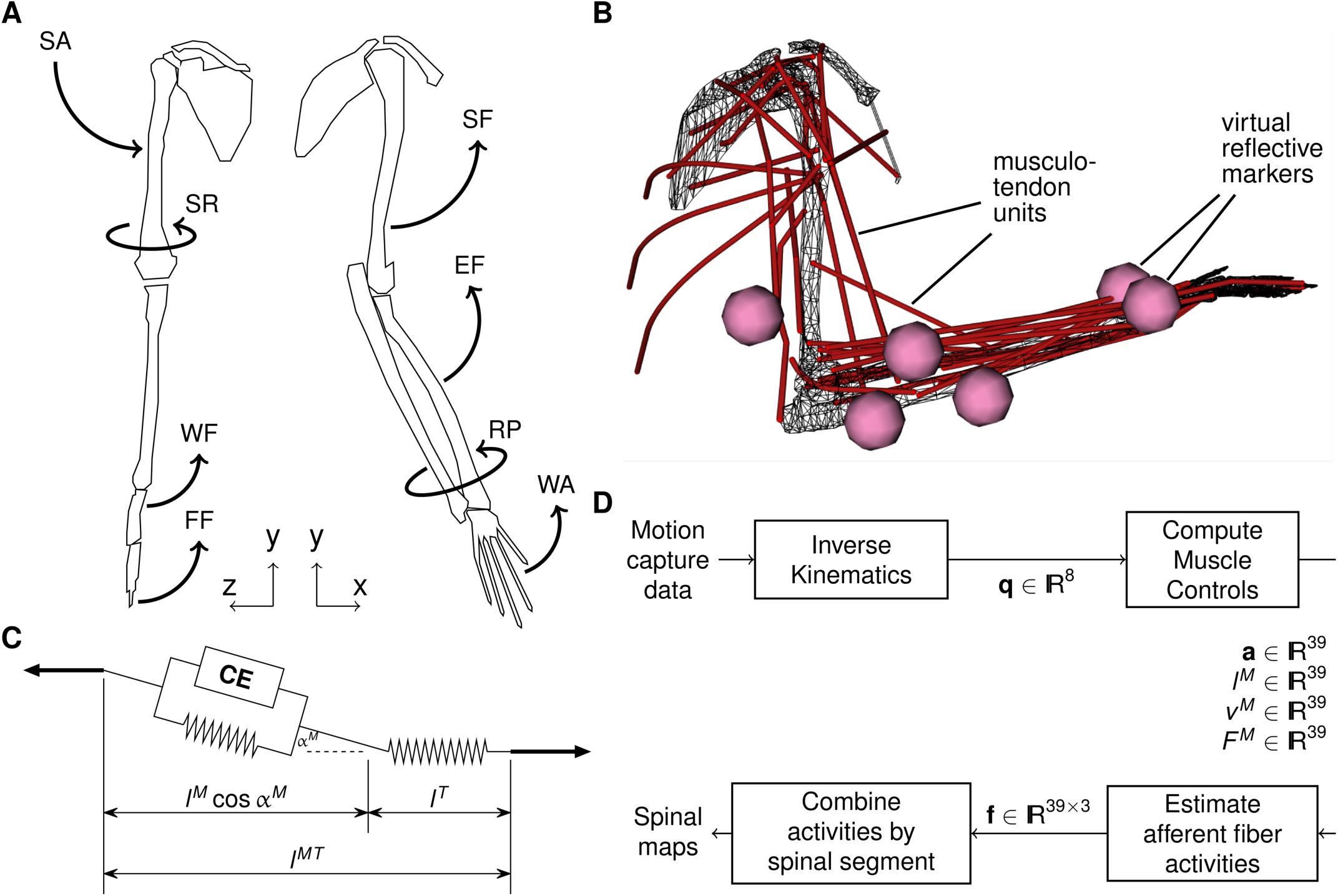
Modelling approach. **A**: the *Macaca Fascicularis* right arm model of 8 bone structures, articulated around 8 degrees of freedom (SA: shoulder adduction, SR: shoulder rotation, SF: shoulder flexion, EF: elbow flexion, RP: radial pronation, WF: wrist flexion, WA: wrist abduction, FF: fingers flexion) **B**: 39 musculo-tendon units (MTU) allow dynamic activation of the joints. 6 virtual markers are added to the model, conform to the placement of real markers on the recorded animal. **C**: in the Hill muscle model, a MTU consists of a Contractile Element (CE) mounted in parallel with a passive element together representing the fiber, mounted in series with a passive element representing the tendon. **D**: Computational flowchart: the joint angles **q** are produced by OpenSim’s inverse kinematics, and are fed to OpenSim’s CMC. The latter yields fiber properties such as the activity **a**, the fiber length *l*^*M*^ and its first derivative *v*^*M*^, as well as the fiber force *F*^*M*^. With linear models developed by Prochazka [34,35], these properties are used to compute 3 types of sensory feedback **f** for each of the 39 MTUs, sensory feedbacks which are then separately mapped to spinal segments to obtain spinal maps.

### A. Biomechanical model

The right arm model includes 39 musculo-tendon units (MTU), 8 bone structures, and 8 joints. We adapted a SIMM (Motion Analysis Corporation, USA) model of the right arm of the *Macaca mulatta* [32] to OpenSim (National Center for Simulation in Rehabilitation Research, USA) and scaled each bone separately to the dimensions of the *Macaca Fascicularis* arm. The parametrization of each arm segment was complemented with mass [36], and resulting inertia matrix coefficients calculated for each segment taken as a homogeneous cylinder. We obtained further anatomical measurements by dissecting an arm specimen of a female *Macaca Fascicularis*. During the dissection, approximate muscle fiber and tendon lengths were also measured. From a dissected muscle, and after removing the tendons, we measured the fiber volume by submerging it in a graduated beaker. We estimated the fiber principal cross-sectional area (PCSA) as the fiber volume divided by the fiber length. Subsequently, the maximal isometric force was estimated as 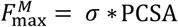, with *σ* = 0.3 [Nm^−2^] [37]. We repeated such measurements for a total of 36 muscles of the arm, hand and shoulder. By combining this dataset with reported morphological measurements of macaques [36, 38, 39], we complemented the model with novel data and adapted it to the *Fascicularis* anatomy. However given the difficulty to measure it, we kept the pennation angle parameter at null [36]. Measurements from a subset of representative muscles is reported in Table 1. Moreover, following the observations made during the dissection, we adapted several MTU lines of action, and added wrapping surfaces when necessary to prevent MTUs from crossing bones. These adjustments preserved the operating ranges of normalized MTU length and moment arm. Finally, we added a joint to improve representation of the hand. The hand bone structure was split into two pieces around the first knuckles, in order to obtain the “fingers” and the “palm” (with the thumb). For finger actuators such as the flexor digitorum superficialis muscle (FDS), the model already included a single MTU whose distal attachment point was located on the palm. However since we wanted to allow for the simulation of power grasps, we made adaptations to the FDS and its antagonist MTU, the extensor digitorum (EDC), so that together they could actuate the new finger joint. Both MTUs were stretched and the tendon lengths increased accordingly, in order to reach opposite sides of the middle finger’s distal phalanx. A degree of freedom was created to allow fingers flexion in the range (−10, 90) degrees, where the flat hand was taken to be the neutral fingers flexion. The complete model is available as supplementary material to this manuscript.

**Table 1:**
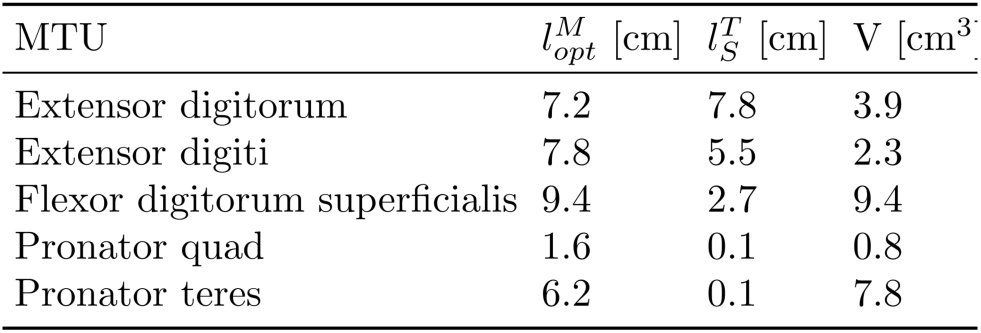
Morphometric measurements of a subset of representative arm and forearm musculo-tendon units (MTU): optimal fiber length 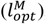, tendon slack length 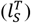, and fiber volume (V).

### B. Afferent fiber model

The average firing rate of group Ia, Ib and II afferent fibers can be estimated from the state of a single MTU at time *t* using equations developed to fit experimental recordings of afferent firing rates in cats by Prochazka and colleagues [18, 34, 35]. MTU sizes are comparable between the cat hind limb and the *Macaca Fascicularis* upper limb, therefore we expect such models to offer a reasonable approximation of sensory fiber dynamics in the *Fascicularis* arm. Specifically, for a given MTU, we approximated the firing rate *f*_*ia*_ of Ia afferents as:

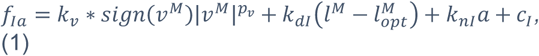

which is the sum of terms that depend on fiber contraction velocity *v*^*M*^ [mm/s], fiber stretch (obtained as the difference between fiber length *l*^*M*^ [mm] and optimal fiber length 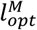 [mm]), the normalized muscle activity *a*, and a baseline firing rate *c*_*I*_. All constants *k*_*_ and *c*_*_, as well as *p*_*v*_, are numerical coefficients that have been previously determined [34, 35].

The firing rate of Ib afferents was estimated to be proportional to the ratio of the force exerted by the muscle fiber *F*^*M*^ [N] over the maximal isometric force 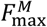[N]:

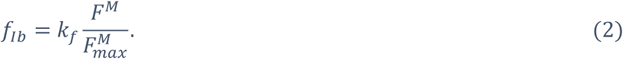

Finally, the firing of group-II spindle afferents was estimated as the sum of terms depending on fiber stretch, muscle activity, and a baseline firing:

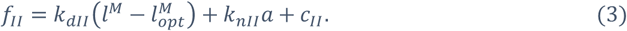

We enforced a lower bound firing rate of 0 [Hz] or [impulses/s] to each fiber population.

### C. Experimental dataset: kinematics

Two females *Macaca Fascicularis* (Mk-Sa, age 7 years, weight 4 kg and Mk-Br, age 4 years, weight 3.5 kg) were trained to reach with the left arm for a spherical object, grasp it, and pull it towards a return position to receive a food reward. Animals were housed within a group of five animals at the University of Fribourg, Switzerland. All experimental procedures were performed at the University of Fribourg in agreement with the veterinary cantonal office of the Canton of Fribourg according to the license n°2017_04_FR.

We recorded three-dimensional arm-joint kinematics using the VICON Vero system (VICON, Oxford, U.K.) with 12 infrared cameras, 6 reflective markers attached to the arm joints, and 2 high definition video cameras. Two sets of n=9 (Mk-Sa), and n=19 (Mk-Br) reaching and grasping successful trials, each cut between the cue command and the return to start position, were extracted and used for the results of this study. Kinematic and video recordings were synchronized and sampled at 100Hz. The recordings of the reflective markers’ positions in 3D were then resampled over 1000 time points. Given the constancy of the trial durations (1.62 ± 0.26 s for Mk-Sa, 2.24 ± 0.20 s for Mk-Br), we proceeded to average in normalized time the marker positions across trials. The duration of this time-normalized average trial was finally scaled back to the real average trial duration in order to be fed to the chain of computations. The time point corresponding to the grasping event was manually identified in each video recording separately. The location of the markers on the arm is shown in Fig. 1. Markers were placed at the middle of the upper arm, at the distal end of the humerus, at the elbow joint and at the proximal end of the ulna, and at the distal ends of the ulna and radius at the wrist. Finally, we artificially triggered a whole hand flexion of the model’s “fingers” upon initiation of grasping by the animal. We simulated the fingers’ flexions by making the fingers’ joint angle follow a logistic function of time fitted to match the start and end angle values, with its step centered on the time point identified as the grasping onset. The key criterion in choosing the logistic function for this artificial joint evolution is that its first derivative is bell-shaped, which is the natural temporal profile of joint velocities [40].

### D. Experimental dataset: electromyography

The monkeys were implanted with chronic bipolar teflon coated stainless steel wire electrodes in the deltoid (DEL), biceps (BIC), triceps (TRI), FDS and EDC muscles of the left arm (Cooner wires). The surgical procedures have been described elsewhere [13]. We recorded differential EMG signals at 12 kHz using a TDT RZ2 system with a PZ5 pre-amplifier (Tucker Davis Technology, USA) and synchronized them with the 3D kinematic recordings using analog triggers. EMG recordings were high-pass filtered at 5Hz, rectified, and low-pass filtered at 6Hz, to obtain signal envelopes for model validation purposes. EMG signal envelopes were normalized in amplitude (divided by their maximal value over the trial) independently for each muscle, and their time course was scaled similarly to that of the kinematic recordings.

### E. Estimation of Spatiotemporal maps

Proprioceptive sensory afferents, innervating muscles and tendons, converge towards the spinal cord in peripheral nerves bundled with their homonymous muscle motor axons. Therefore, we assumed that their organization within the dorsal roots matches that of their homonymous motor axons in the corresponding ventral roots. Following this assumption, the afferent activity stemming from each MTU was mapped to the dorsal roots and thus to the spinal segments using the rostro-caudal distribution of motor pools in the primate spinal cord [41].

As data were missing for the deltoid in the mentioned publication, we approximated its motor pool localization using data available in humans [42]. Table 2 shows the resulting proportions of motoneurons of each muscle in each cervical spinal segment, which we assumed to represent the proportions of afferent fibers of each muscle projecting to each segment as well.

**Table 2:** Proportional distribution of the motor pools of deltoid (DEL), biceps (BIC), triceps (TRI), flexor digitorium superficialis (FDS) and extensor digitorium communis (EDC).

| Spinal segment | DEL | BIC | TRI | FDS | EDC |
| --- | --- | --- | --- | --- | --- |
| C5 | 0.20 | 0.20 | 0.01 | 0.00 | 0.00 |
| C6 | 0.47 | 0.47 | 0.05 | 0.00 | 0.00 |
| C7 | 0.25 | 0.25 | 0.26 | 0.00 | 0.03 |
| C8 | 0.08 | 0.08 | 0.53 | 0.26 | 0.65 |
| T1 | 0.00 | 0.00 | 0.15 | 0.73 | 0.32 |

We estimated the input sensory activity *a*_*i,x*_ of type *x*, received by the *i-th* spinal segment, as:

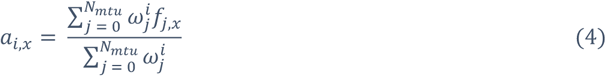

where *f*_*j,x*_ is the firing rate of the proprioceptive fibers of type *x*of the *j-th* MTU, and 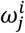 the proportion of its afferent fibers projecting to the *i-th* spinal segment. The simulated and recorded data. In particular, simulated joint angle trajectories are well within the experimental variability range of recorded data (R=0.90 for Mk-Sa and R=0.81 for Mk-Br). We then compared the computed muscle activities and the envelopes of recorded EMG signals from upper limb muscles. Qualitative analysis of activity patterns shows that the simulated muscle activities match the recorded EMGs (Fig. 3). In particular, upper arm muscles are activated in the first part of the reaching phase to lift the arm and initiate the whole limb movement. Successively, forearm and hand muscles are activated to shape the grasp, and grab the object. Finally, biceps and deltoid muscles are strongly activated during the pulling phase of the movement. Quantitative comparison between simulated muscle activities and EMG envelopes shows good correlation levels for almost all muscles in both animals (Fig. 3). Results considerations that led to the expression of *a*_*i,x*_ are addressed in the discussion.

**Fig. 2:**
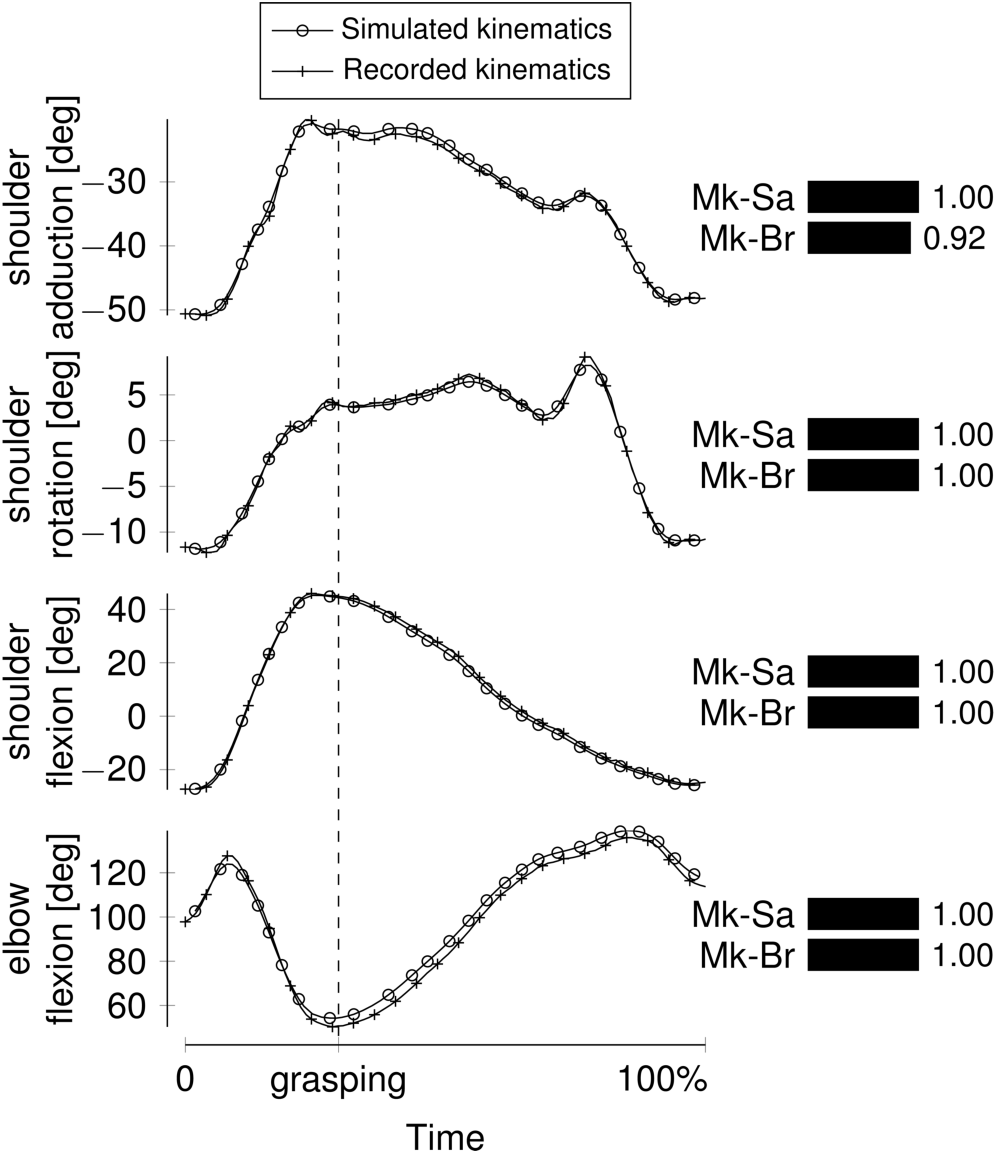
Evolution of the principal joint angles during the three-dimensional reaching and grasping task for Mk-Sa, recorded and simulated (resulting from the estimated muscle activities). The time of grasping was manually identified for each recording separately. Joint angles computed by forward dynamics, using estimated muscle activities, are in excellent agreement with experimental recordings as can be quantified with the cross-correlation between the two curves, presented in bars.

**Fig. 3:**
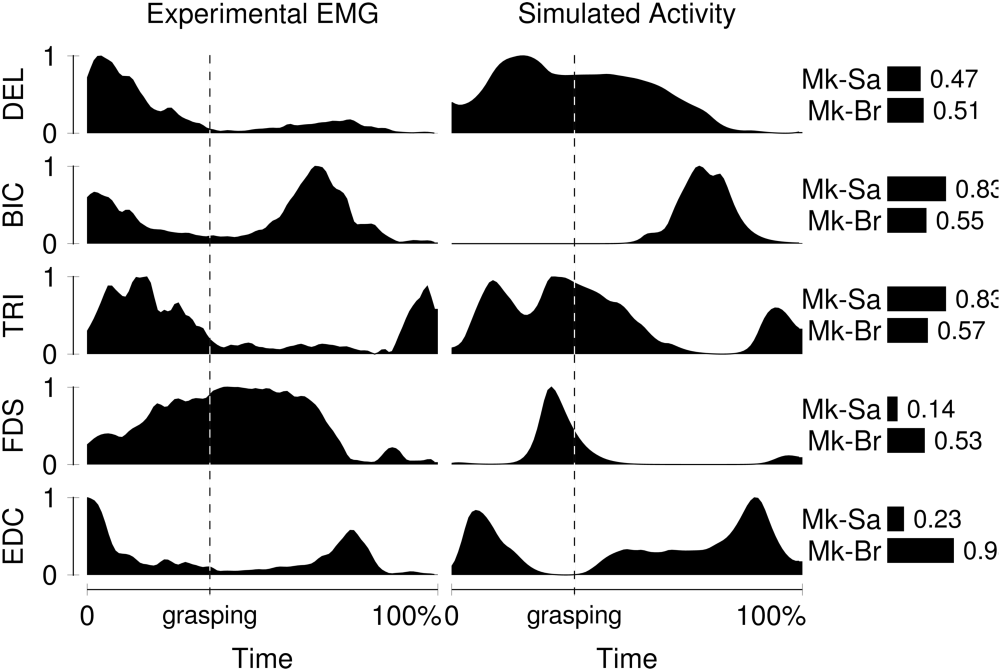
Averaged and normalized EMG envelopes, compared to computed muscle activity for Mk-Br. DEL: deltoid, BIC: biceps, TRI: triceps, FDS: flexor digitorum superficialis, EDC: extensor digitorum. Correlation between the two curves, for each monkey, are shown in bars.

The set of *a*_*i,x*_ ‘s can thus be represented as a color image summarizing the amount of input sensory activity of type *x* received by the cervical spinal cord over time, similarly to spatiotemporal maps of motoneuronal activity [12, 43].

## III. RESULTS

### A. Model Validation

We recorded simultaneous 3D kinematics of the upper limb and EMGs of the principal upper limb muscles during an unconstrained three-dimensional reaching, grasping and pulling movement. To validate our biomechanical model, we fed averaged 3D trajectories of the joint markers to OpenSim, and computed joint angles with inverse kinematics. We then used the compute muscle control (CMC) tool to estimate a set of muscle activities, that represented a plausible solution to the inverse biomechanical problem, i.e.: what is the set of muscle activities, from which the recorded motion of the arm has originated? Next we fed the simulated muscle activities to OpenSim’s forward dynamics, thereby obtaining simulated kinematics solution to the forward biomechanical problem. Comparing the kinematics produced with this approach against the experimental joint angles (Fig. 2) shows excellent similarity between corresponding to cross-correlation values of about 0.50 correspond to model predictions that seem reasonably accurate. While the predictions realized for the finger actuators of Mk-Sa fall short of this accuracy, we observe outstanding accuracy for the elbow actuators of Mk-Sa, and for finger actuators of Mk-Br.

### B. Spatiotemporal map of motoneuronal activity

We then projected the computed muscle activity onto the spatial locations of arm motoneurons in the primate cervical spinal cord (Table 2, Fig. 4). Clear bursts of motoneuronal activity span the cervical spinal cord from rostral (C5-C6) to caudal segments (C8-T1) reflecting the upper-arm to forearm to finger muscles sequence of activation. Strong bursts of motoneuronal activity are present in rostral segments at the beginning of the reaching phase, whereas more caudal segments, where motoneurons of intrinsic and extrinsic hand muscles are located, are activated around the grasping phase. Finally, rostral segments are again activated but with a lower amplitude during the pulling phase. These results are in agreement with experimental findings in human subjects performing a similar task [44].

**Fig. 4:**
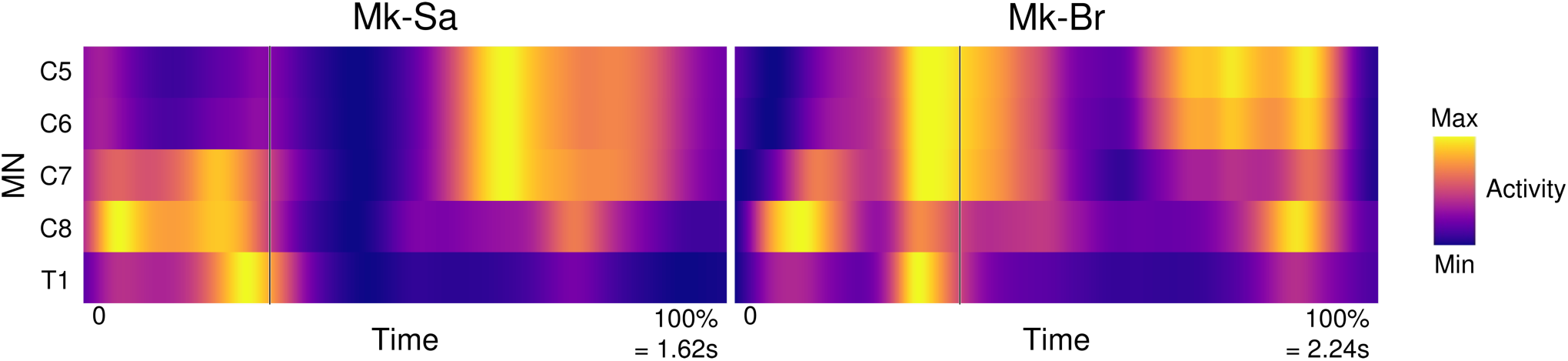
Simulated spatiotemporal map of motoneuronal activity during the three-dimensional reaching task for both monkeys, obtained using the weights presented in Table 2. Normalized activity over time and for each segment is shown in a color scale from purple (no activity) to yellow (maximal activity).

### C. Sensory Afferent Firing rates

We estimated the firing rates of group Ia, Ib and group II afferent fibers by feeding the simulated muscle stretches and forces to the mathematical model of afferents firing rates (1), (2), (3). Our framework allows the direct comparison between the firing rate of each simulated sensory fiber ensemble (consisting of the fibers of a specific type originating from a specific MTU) and its homonymous muscle activity during a whole limb three-dimensional movement (Fig. 5). For simplicity we reported here the example of the agonist /antagonist of the elbow, i.e. biceps and triceps. As expected from intuition, antagonist Ia afferents are anticorrelated. Ia afferents of the biceps are most active when the triceps’ are least active and vice versa. Instead, Group II and Ib afferents are not entirely anti-correlated between these antagonists. Moreover, in the case of the triceps, afferents show anticorrelation with active muscle contraction. This is surprising considering that group II and Ib afferents should respond more to active muscle force. Likely this is the result of both passive and active tendon elongation and muscle stretches that occur during multi-joint movements.

**Fig. 5:**
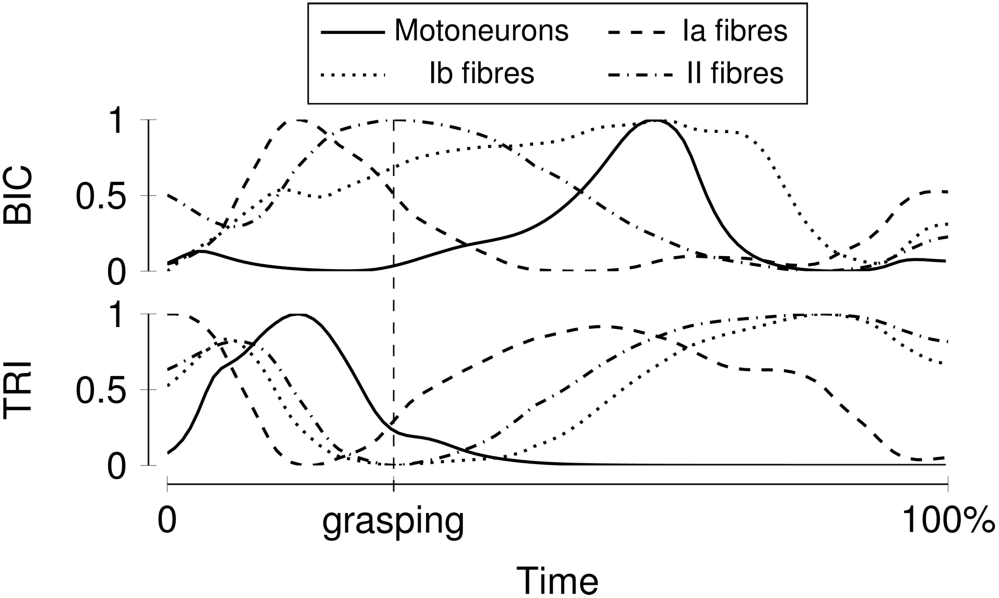
Profiles of normalized moto- and sensory-neurons firings, for the principal arm and forearm muscles in Mk-Sa. Motoneuron firing rates are shown with solid lines, Ia dashed, II dashed-dotted and Ib dotted.

### D. Spatiotemporal map of proprioceptive inputs

We then projected the activity of the proprioceptive sensory afferents onto the spinal segments. Individual sensory input maps for the Ia, Ib and group II fibers, are reported in Fig. 6.

**Fig. 6:**
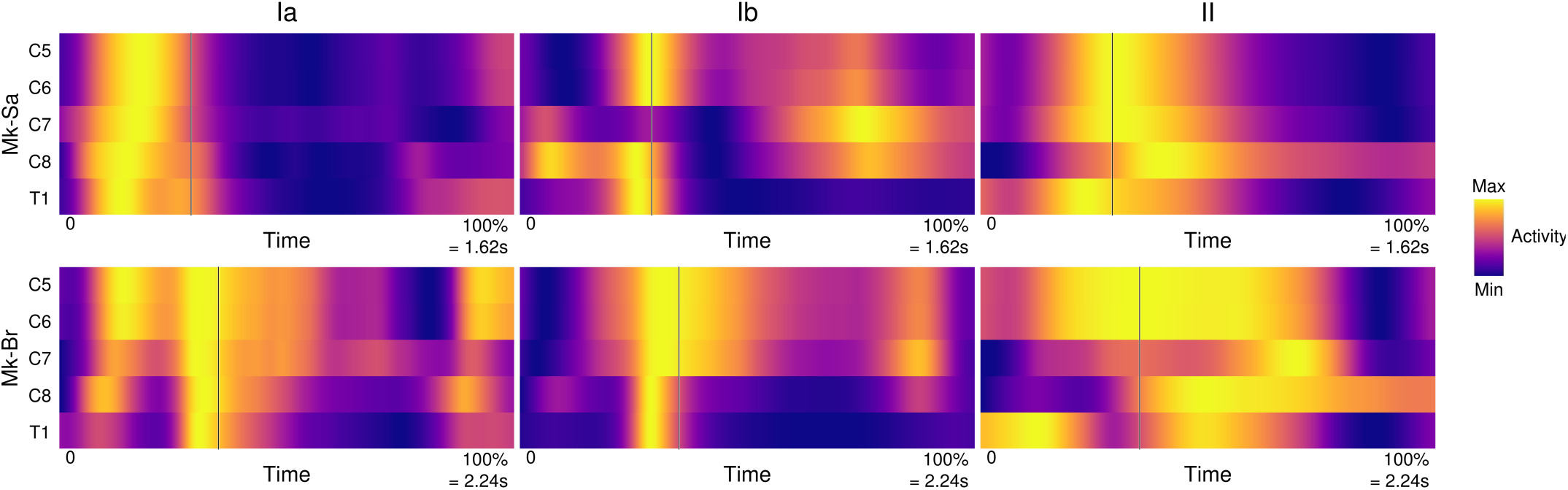
Spinal map of the three different sensory feedbacks during a standardized reaching and grasping task, for both animals. Here as well, the three identified phases present noticeably different patterns of afferent activity.

These maps show very distinct patterns in both space and time for each of the fiber populations across movement in both subjects. While Ia activity precedes the grasping phase, Ib, and group II activity is maximal during the pulling phase when the animal applies maximal force. However, Ib activity shows sharp activations along the whole cervical enlargement while group II has long bursts of activity that span the entire duration of the pulling phase. Finally, we computed the total normalized proprioceptive sensory activity received by the cervical spinal cord by summing the normalized activities of each fiber type. The resulting spatiotemporal map (Fig. 7) shows how total proprioceptive inputs are conveyed in space and time to the cervical spinal cord during three-dimensional reaching movements. Overall this map is qualitatively similar to the spatiotemporal map of motoneuronal activity (Fig. 4). Proprioceptive activity first arises in the rostral segments, moves towards the caudal segments during grasp pre-shaping and finally peaks in both rostral and caudal segments during the pulling phase. In particular, it is sustained for the entire duration of the motor bursts responsible for the movement execution.

**Fig. 7:**
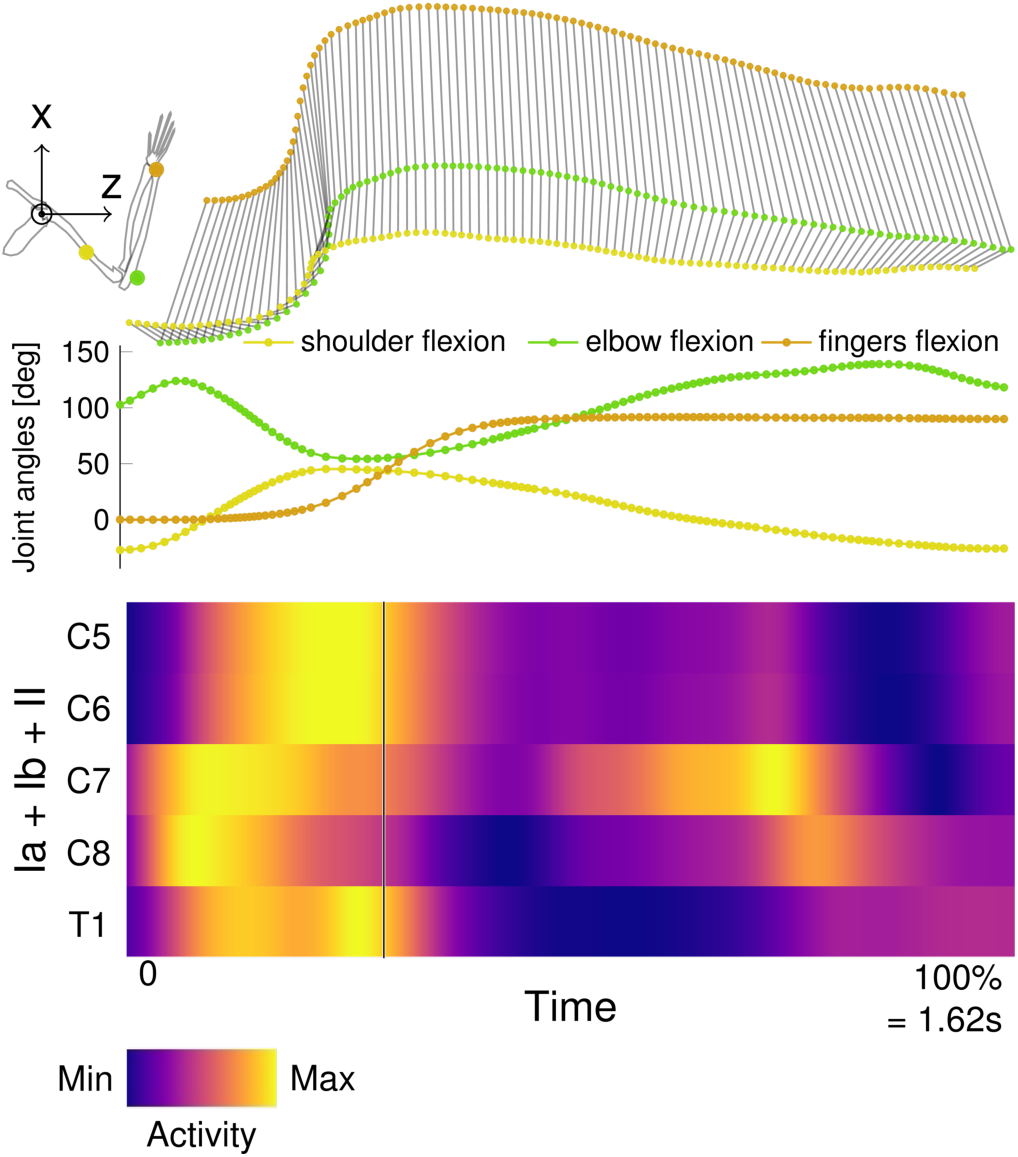
Spinal map of combined estimated proprioceptive feedbacks of Mk-Sa, during a standardized reaching and grasping task. The three identified phases present noticeably different patterns of general afferent activity.

## IV. DISCUSSION

We extended and validated a biomechanical model of the arm of *Macaca Fascicularis* to predict the firing rates of ensembles of proprioceptive afferents during three-dimensional reaching and grasping.

### A. A realistic primate arm model

We reworked a model of the rhesus monkey upper limb to fit the geometrical and mechanical properties of the *Macaca Fascicularis* arm. We dissected most of the muscles of the arm, forearm and shoulder from a *Fascicularis* arm specimen and extracted parameters such as fiber and tendon lengths, as well as fiber volume for each of the analyzed muscles, in order to refine and complement the initial model parametrization. As was the case in the original model, the joint angle space was constrained to physiological values.

Our model was able to faithfully reproduce the recorded kinematics, thus validating the skeletal model and the muscular parametrization as a whole. When comparing the computed muscle activity with the envelopes of the recorded EMG signals, we found a weaker correspondence. However, quantitative discrepancies between simulated and recorded muscle activities are common in biomechanical models [45, 46]. Specifically, since we compute muscle activities from joint kinematics our model cannot simulate co-contraction of antagonist muscles (e.g. as occurs during stiffening of the arm). This limited the quality of forearm EMG estimation and will be the object of future research directions. Such discrepancies could furthermore originate in the optimization strategy chosen to compute the solution of the inverse biomechanical problem. There is indeed redundancy in muscle space, that requires imposing a strategy to extract one set of muscle activities, amongst several that are able to produce a given motion. In the case of OpenSim, the choice is to minimize the sum of squared MTU activities, i.e. minimizing metabolic energy consumption [47]. Other strategies might be explored to improve future studies results. Finally, we did not expect full correspondence between estimated and recorded muscle activity because the model parametrization does not take into account the subject-specific detailed anatomy [48, 49] and physical training [50, 51]. Nevertheless, our simulated muscle activity dynamics and kinematics are overall similar to those yielded by other investigations involving non-human primate upper limb models [32], strengthening our confidence in the validity of our approach [46, 52].

Finally, it may be worth noting that the biomechanical model presented here can be embedded in a broader closed-loop simulation environment. The estimated sensory activity could be used to compute motoneuronal activity, itself could be driving the evolution of arm kinematics, and in turn allow to estimate an updated distribution of sensory activity. Such an approach may enable testing for different hypotheses regarding the sensorimotor control of the upper limb.

### B. Sensory afferent firing dynamics

The firing dynamics of primary sensory afferents during active functional movements is a key information to study sensorimotor integration during voluntary movement execution. Additionally, modern neuroprosthetic applications aiming at the recovery of both motor [24] and sensory [23, 53] functions in patients affected by neurological disorders often seek to design biomimetic stimulation protocols, with the underlying assumption that the most effective therapy depends on reproducing the natural activity in primary sensory afferents as closely as possible. For both these basic and translational applications, knowledge about the firing dynamics of various types of afferent fibers during functional voluntary movements is required. A sufficiently accurate computational model can be used to estimate these firing rates during multi-joint movements in dynamic tasks. Such estimates can support the interpretation of experimental data, as well as assist the design of neuroprosthetic systems that aim at reproducing these firing rates. Towards this goal, we describe a method to study sensory fiber ensembles from multiple muscles simultaneously during voluntary movements. The results that we reported show the importance of studying these signals during functional tasks. Indeed, when looking at the Ib afferent firing rates, we notice that in the triceps, the Ib afferents are anti-correlated with muscle activation (Fig. 5). Ib afferents encode force information via tendon elongation and are thus commonly expected to be active during muscle contraction and consequent tendon elongation. However, during a multi-joint movement, active and passive tendon elongation can also occur due to the contraction of antagonist muscles, or gravity compensation. Therefore, large discrepancies from the expected firing patterns of this fiber population may emerge as a result of complex bio-mechanical interactions, as it was likely the case in our simulations.

### C. Spatiotemporal patterns of proprioceptive input to the cervical spinal cord

We assumed that the proprioceptive afferents are distributed along the rostrocaudal extent of the cervical spinal cord similarly to their homonymous motoneurons. Given the well-known strong monosynaptic connectivity between muscle spindle Ia afferents and motoneurons, this approximation seems reasonable [54]. Using this assumption, we estimated the spatiotemporal distribution of the proprioceptive input to the spinal cord during arm movement. The total proprioceptive activity reaching the spinal cord during movement is a combination of both spindle and Golgi tendon fibers activity. This activity (Fig. 7) arises in the form of clear bursts that span the spinal segments and are sustained across the entire duration of movement whilst being strongly modulated. This is in agreement with the well-known fact that the spinal cord receives large amounts of neural inputs during movement, and that spinal circuits are continuously fed with information. Moreover, the neural input supplied by the different sensory fiber ensembles present markedly distinct spatiotemporal patterns, suggesting that a stimulation-based restoration of “proprioceptive” information and perception must target these three fiber populations independently.

### D. Insights for the design of neuroprosthetic systems

Our results offer important insights for at least two applications in neuroprosthetics. The first important observation regards the distinct spatiotemporal patterns of each specific fiber population. Modern biomimetic strategies that aim at restoring sensation [53] employ electrical stimulation of the peripheral nerve to convey information to the central nervous system of amputees. However, this technology does not allow selectivity on fiber types [53], This is particularly true for Ia and Ib fibers. Indeed, these afferents have similar diameters and thus similar recruitment thresholds making it challenging to independently control their firing rates. These fibers convey complementary information about movement and force, and our simulations show that they are active at markedly different moments during movement execution. This poses important questions on the theoretical limitations of electrical stimulation technologies to achieve realistic proprioceptive feedback in amputees.

The second consideration concerns technologies aiming at the stimulation of the spinal roots such as EES. When active, electrical stimulation of a specific root will cancel the natural flow of information of each recruited afferent and substitute it with the imposed stimulation frequency [24]. However spinal circuits require correct flow of sensory feedback to be able to produce functional movements. Therefore, development of epidural stimulation strategies of the spinal cord must take into account the spatiotemporal maps reported in Fig. 7. For instance, the T1 spinal roots is supposed to have no input activity both at the beginning of reaching and at the end of the pulling phase. This means that stimulation targeting that root in these periods should be avoided to prevent delivery of aberrant proprioceptive information. Similar consideration can be made for the other roots.

### E. Model limitations

Our model is limited by the data available for primates. Morphometric measurements, as well as live recordings, are scarce and scattered. The number of MTUs studied to obtain the spatiotemporal maps should be extended when data regarding motor pool distributions of additional muscles will be made available. Hence the reported spatiotemporal maps of proprioceptive input are built using a limited set of arm and forearm muscles. Yet, the actual proprioceptive input received by a spinal segment is the number of action potentials per unit time entering that spinal segment via all the proprioceptive fibers running in the corresponding dorsal root. However, the exact distribution of proprioceptive fiber ensembles in the dorsal roots remains to date surprisingly unknown. We thus limited our analysis to the 5 MTUs shown in Fig. 3, for which we could estimate the relative proportions of fibers in the different spinal roots. We assigned identical weights to the Ia-afferent pools of every represented MTU (see *a*_*i,x*_ (4)). This is equivalent to assuming that similar stretches and applied forces in these MTUs induce equal amounts of proprioceptive input to the spinal cord. This assumption may not hold if large differences in the absolute number of proprioceptive fibers exist between MTUs. The normalization introduced in (4) eases the interpretation of the spatiotemporal maps. Without this term, the input sensory activity estimated using (4) would be biased towards those segments for which 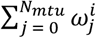 is larger (e.g. C6 compared to C5) and the temporal variations in the unfavored segments would have been obscured. In summary, the spatiotemporal maps shown in Fig. 6 and 7 are best interpreted in terms of normalized temporal variations of the combined proprioceptive input emerging from the selected MTUs and received by individual segments, rather than actual amounts of neural input expressed in impulses/sec.

## CONCLUSION

We presented a computational estimation of the spatiotemporal patterns of proprioceptive sensory afferent activity during three-dimensional arm movements in a clinically relevant animal model. We showed that the patterns of proprioceptive inputs during functional movements are surprisingly complex and do not necessarily match intuition. Additionally, we showed that different fiber populations have markedly distinct spatiotemporal patterns of activity, highlighting the need of recruiting these populations independently to restore the natural flow sensory information. Finally, our model can be integrated in a broader in-silico platform to simulate the effect of electrical stimulation of the sensory afferents on arm biomechanics, as well as support basic studies on sensory systems. These advancements can thus reduce the number of animals involved in invasive experiments.

## Supporting information

Arm_Model

## ACKNOWLEDGMENTS

We thank Prof. Eric M. Rouiller for supervising the animal experiment protocols, providing anatomical specimens, and for his continuous support and insightful scientific discussions. We thank Prof. Grégoire Courtine and Prof. Jocelyne Bloch for implanting the EMG leads. Mrs Maude Delacombaz for carefully training the animals. Finally, we thank Mr. Jacques Maillard and Mr. Laurent Bossy for their meticulous work in the care of the animals. This work was supported by a Swiss National Science Foundation Ambizione Fellowship to M.C., a grant from the Wyss Center in Geneva (WCP 008) and a Catalyst Fund grant from the Bertarelli Foundation N8C1709 to M.C.

